# Genetic diversity of tomato brown rugose fruit virus in Morocco

**DOI:** 10.64898/2026.05.11.724243

**Authors:** Ayoub Maachi, Livia Donaire, Miguel A. Aranda

## Abstract

Tomato brown rugose fruit virus (*Tobamovirus fructirugosum*) is an emerging virus that affects tomatoes, capsicum, and chili. Since its first detection in Jordan in 2015, the virus was reported in more than 40 countries across all the continents. In Morocco, the virus was reported for the first time in October 2021. However, its genetic diversity remains unexplored. In this work, we used a collection of tomato fruits from local markets to investigate the variability of the virus in the country. We explored the different pressures acting on the N-terminus of the RNA-dependent RNA polymerase, the movement protein, and the coat protein genes. Then, we used haplotype network analyses to reveal the population structure within the Moroccan isolates and studied their relationships with the ones from the world. We found that genetic diversity is low, which is consistent with the global situation. No signatures of diversifying selection were detected across the analyzed genes. However, the virus sequences from Morocco showed a clear geographic structure, suggesting that geographic factors probably combined with agricultural practices may contribute to shaping the population structure of ToBRFV in Morocco.

## Introduction

Tomato (S*olanum lycopersicum L*.) is one of the most important vegetable crops in Morocco, with an estimated production of 1,686,615 tons in 2024 (FAOSTAT; http://www.fao.org/faostat/; accessed in March 2026). Tomato cultivation is very intensive and concentrated mainly in the Souss-Massa region, with significant acreage devoted exclusively to greenhouse production. Viral diseases are considered the major threats to tomato intensive cultivation, which are responsible for significant yield and fruit quality losses (Jones, 2021). The main viruses frequently reported to affect tomatoes are tomato yellow leaf curl virus and pepino mosaic virus (Rivarez et al., 2021). Nevertheless, emerging viruses, i.e., those that have recently appeared in a population for the first time, or that have existed previously but are rapidly increasing in incidence or geographical range, often cause the most important problems. A recent example of an emerging virus infecting tomato crops is tomato brown rugose fruit virus (ToBRFV) (*Tobamovirus fructirugosum;* family *Virgaviridae*).

ToBRFV is an emerging tobamovirus that affects tomato, capsicum, and chili (Salem et al., 2023; Zhang et al., 2022; Caruso et al., 2022). In tomato and depending on the variety, ToBRFV may cause leaf chlorosis, mosaic patterns, and mottling on younger leaves. On the fruits, the virus may cause chlorotic spots and marbling, deformation and uneven ripening, and brown rugose patches (Caruso et al., 2022). ToBRFV is a stable and highly infectious virus that is easily transmitted by mechanical means, via seeds (Davino et al., 2020; Salem et al., 2022), bumblebees (Levitzky et al., 2019), and via contaminated water (Mehle et al., 2023) and soil (Molad et al., 2024), enabling its spread locally and over long distance. The virus can persist for a long period of time on different structures in production sites (Skelton et al., 2023; Ehlers et al., 2023). Since it is first report in Jordan in 2015, the virus was reported in more than 40 countries across all the continents. ToBRFV showed exponential population growth until 2020, followed by a decline in effective population size, likely due to control measures or saturation (Ibañez et al., 2025). The variability of ToBRFV is limited, with countries experiencing single or multiple introduction events (Ibañez et al., 2025). In Morocco, ToBRFV was first reported in October 2021 in the Souss-Massa region and in March 2022 in the Dakhla region (EPPO Reporting Service: 2023/2025).

In this study, we aimed to investigate the genetic diversity of ToBRFV from Morocco. To do so, we used a collection of tomato fruits from local markets. We investigated the variability across a genome fragment comprising the N-terminus of the RNA-dependent RNA polymerase (RdRp), the movement protein (MP), and the coat protein (CP) genes, then explored the different pressures acting on each of these genes. We used haplotype networks to reveal the population structure among the Moroccan isolates, and their relationships with the ones worldwide. Our findings suggest limited introductory events to the country with geographic factors contributing to shaping the population structure of ToBRFV from Morocco.

## Material and Methods

### Sample collection and nucleic acid extraction

One hundred tomato fruit samples, exhibiting chlorosis and uneven ripening were collected from shops in markets located near the main tomato producing areas in the Souss-Massa region, except from two sites which were located inside the city of Agadir. Samples were collected from seven different market sites during the period from May to June 2025. In each market, one to three shops were visited (Figure 1A), and four to eight samples were collected from each shop. Total RNA was extracted using the RNA preparation method from Souiri et al., with modifications (Souiri et al., 2017). Briefly, 100 mg of pulp, skin and juice of tomato was ground with 100 μl of sterile-distilled water in a 1.5 ml microtube using cotton swabs, then total volume was made up to 1 ml with sterile-distilled water. After precipitation for 1 h at 4°C, the supernatant was recovered in a new microtube, and RNA was precipitated using 0.1 volume of 3M NaOC (pH 5.2) and 2 volumes of absolute ethanol. The pellet was washed using 1 ml of 70% ethanol and kept at 4°C for downstream analysis.

**Figure 1.**
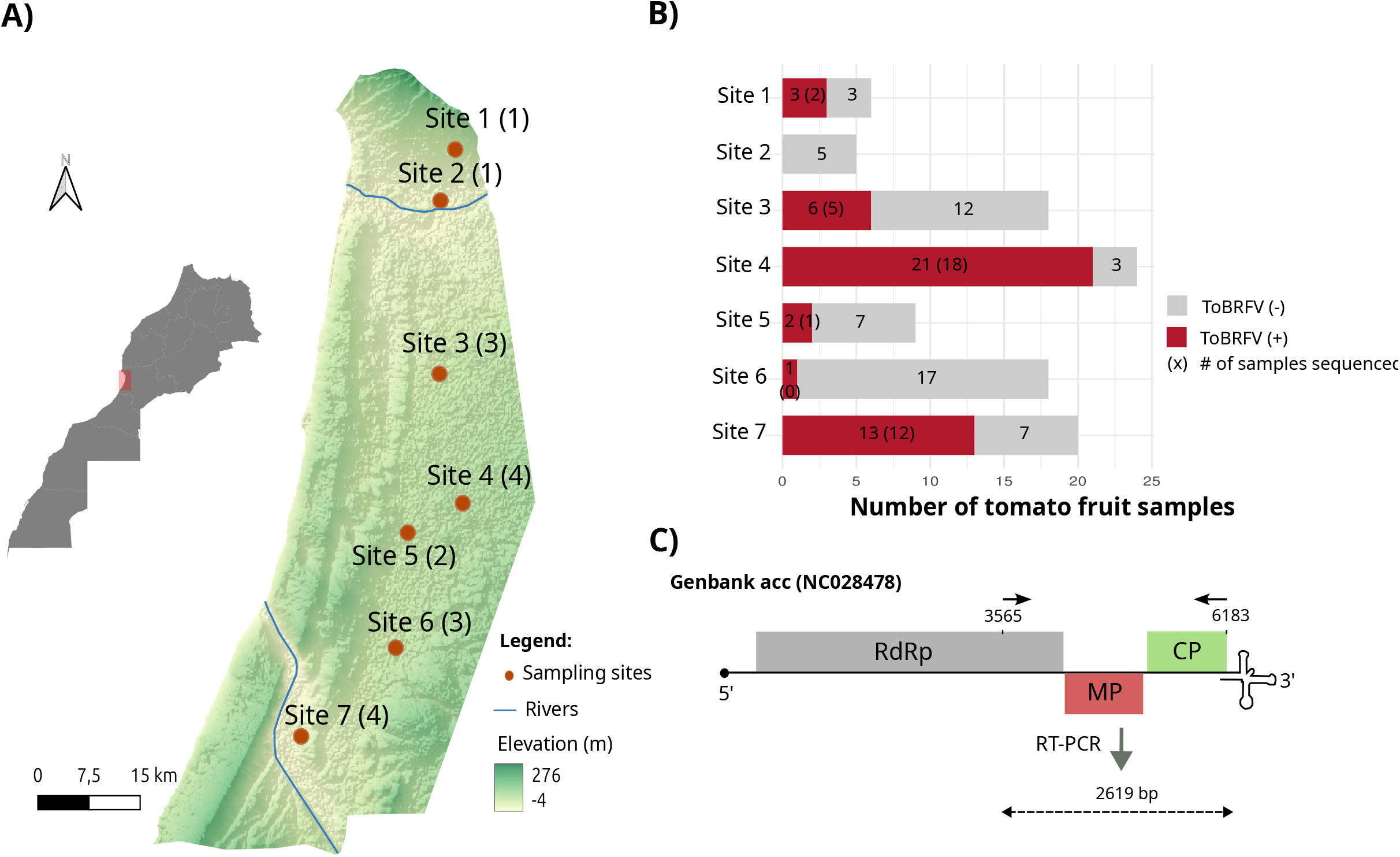
Fruit collection sites and tomato brown rugose fruit virus (ToBRFV) (Tobamovirus fructirugosum) detection from different vegetable markets. (A) Map showing the location of the different markets visited for sample collection. The figures in brackets indicate the number of shops visited per market. (B) Number of tomato fruit samples collected from each site, and the incidence of ToBRFV from each market. The figures in brackets indicate the number of samples sequenced for genetic diversity analysis. (C) Diagram representing the ToBRFV genome. The position of the primers and the genomic region used for virus detection and genetic diversity analyses are indicated.

### ToBRFV detection and amplicon sequencing

ToBRFV was detected in individual fruit samples by RT-PCR using primers LD32F: 5’-TGTTACCATGAGGTTGACTGAC-3’ and LD35R: 5’-GGTGCAGAGGACCATTGTAA-3’ designed for this study (Figure 1C). The specificity of the primers was confirmed *in silico* by a blastn search against the NCBI database. Complementary DNA was synthesized with SuperScript III reverse transcriptase (Thermo Fisher Scientific, Waltham, USA) using the primer LD35R. Each 20 μl reaction consisted of 0.2 µl of 10 µM LD35R, 1 µl of 10 mM dNTP mix, 4 µl of 5x First Strand Buffer, 1 µl of 0.1M DTT, 1 µl of RNase inhibitor, 1 µl of SuperScript III, 8.8 µl of sterile-distilled water and 3µl of RNA.

PCR reactions were carried out using the CloneAmp HiFi (Takara Bio, Kusatsu, Japan). Each 25 µl reaction consisted of 12.5 µl of CloneAmp HiFi PCR premix, 0.5 µl of LD32F/LD35R (10 µM), 2 µl of cDNA, and 9.5 µl of sterile-distilled water. The reactions were incubated in the thermocycler under the following conditions: 98°C for 2 min, 35 cycles (98°C for 10 sec, 55°C for 10 sec, and 72°C for 2 min) and 72°C for 3 min. The PCR products with the expected size (≈ 2,619 bp) were purified using the NZY Gelpure kit (NZYtech, Lisboa, Portugal) and sequenced using amplicon sequencing from Oxford Nanopore Technologies at Eurofins Genomics.

The generated sequences were complemented using 11 additional ToBRFV sequences from GenBank at the National Center for Biotechnology Information (NCBI). These sequences were obtained from tomato fruits in the United Kingdom (UK) and the Netherland (NL), but collected from Morocco between 2023 and 2025 (NCBI). The accession numbers of the complementary data are: PP099972 (NL, 2023), PV389033 (NL, 2024), PV389019 (NL, 2024), PV389020 (NL, 2024), PV389048 (NL, 2025), PV389021 (NL, 2024), and PV389041 (NL, 2024), PV389010 (NL, 2025), PP236969 (UK, 2023), PP236971 (UK, 2023), and PP236970 (UK, 2025).

### Genetic diversity, SNP calling and selection pressures analyses

Multiple sequence alignments were performed using MUSCLE v5 with the -super5 option (Edgard 2022), followed by manual curation. DnaSP6 software was used to calculate the nucleotide diversity (π) in the whole fragment and in individual genes (partial RdRp, MP, and CP) using the Jukes and Cantor method.

Genetic variability was assessed using Shannon entropy. For each alignment position, the Shannon entropy (H) is calculated based on the frequency of each nucleotide, using the following formula *H=-Σx*_*i*_*log*_*2*_*x*_*i*_, where x_i_ represents the frequency of nucleotide i at a given position.

SNP calling was performed on GeneiousPrime, by mapping each of the genes to the ToBRFV reference sequence under the accession number NC028478 and using the “Find Variations/SNPs” function with a minimum variant frequency set to 0.05.

Selective pressures acting on entire genes were determined using BUSTED (Branch-site Unrestricted Statistical Test for Episodic Diversification), while selective pressures acting on individual codons were determined using FEL (Fixed Effects Likelihood) which uses a maximum-likelihood approach to infer nonsynonymous (dN) and synonymous (dS) substitution rates on a per-site basis. Both algorithms were run on the Datamonkey web server (datamonkey.org).

### Haplotype network reconstruction and phylogenetic analyses

Haplotype networks (HN) were constructed using the PopART 1.7 software using the median-joining haplotype network, which is a common method to depict relationships between closely related sequences of the same species or population (Bandelt et al., 1999).

A HN was built using only sequences from Morocco. Another HN was generated by combining ToBRFV sequences from Morocco and the world. 473 sequences of ToBRFV, longer than 6,148 nts, were downloaded from the NCBI Virus database. Sequences were mapped to ToBRFV (1) sequence (PZ370457, generated in this study), and trimmed to comprise the partial RdRp, the MP, and the CP. 270 unique sequences were kept for the analyses using the Seqkit tool. These sequences were merged with the ones generated from this study and were used for the analyses.

### Structural prediction and residue analysis of the MP

527 movement protein sequences were retrieved from the NCBI Virus database (https://www.ncbi.nlm.nih.gov/labs/virus/vssi/#/) using the Virus/Taxonomy: “tomato brown rugose fruit virus; taxid 1761477”. Incomplete sequences were removed and the remaining ones were added to the sequences generated from this study. SeqKit tool was used to remove redundant sequences, yielding 69 unique sequences. Genetic variability within the MP was determined using Shannon entropy and ConSurf (Ashkenazy et al., 2016).

Protein structure of the movement protein (Accession number: WEX498921.1) was predicted using AlphaFold3 (Abramson et al., 2024) via the online server (https://alphafoldserver.com/). Predicted structure was converted from CIF to PDB format using the mCIF-to-PDB converter tool (https://project-gemmi.github.io/wasm/convert/cif2pdb.html). Specific sites were annotated on the 3D structure using PyMOL 3.1 (https://www.pymol.org/).

## Results

### ToBRFV detection in tomato fruit samples

We sampled tomato fruits from eighteen shops located in seven different local markets within the Souss-Massa region (Figure 1A). The region contributes the most to tomato production. The visited markets were located near the tomato cultivation zones. Among the one hundred samples collected, molecular detection by RT-PCR revealed ToBRFV infection in fifty samples. Positive samples ranged from one fruit in Site 6, to twenty-one fruits in site 4 (Figure 1B). No sample from Site 2 was positive (Figure 1B).

### Genetic diversity and selective pressures among ToBRFV sequences

We sequenced a genomic region comprising ≈37% of the ToBRFV genome covering the partial RdRp gene, the MP, and the CP genes (Figure 1C). We sequenced this segment for 38 samples, 1 sequence per infected fruit, and we complemented our data with 11 additional sequences of ToBRFV from tomato fruits from Morocco; these were previously determined and made publicly available in the GenBank database. Pairwise nucleotide identity among the sequences ranged from 99.45% to 100% (Figure 2A). Overall, the nucleotide diversity was low (π = 1.91 × 10^−3^ ± 1.7 × 10^−4^), and diversity was unevenly distributed across the sequenced fragment (Figure 2B) with the RdRp showing the lowest diversity (π = 1.45 × 10^−3^ ± 1.8 × 10^−4^), followed by the MP (π = 1.89 × 10^−3^ ± 10^−7^), and the CP which exhibited the highest diversity (π = 3.08 × 10^−3^ ± 10^−7^).

**Figure 2.**
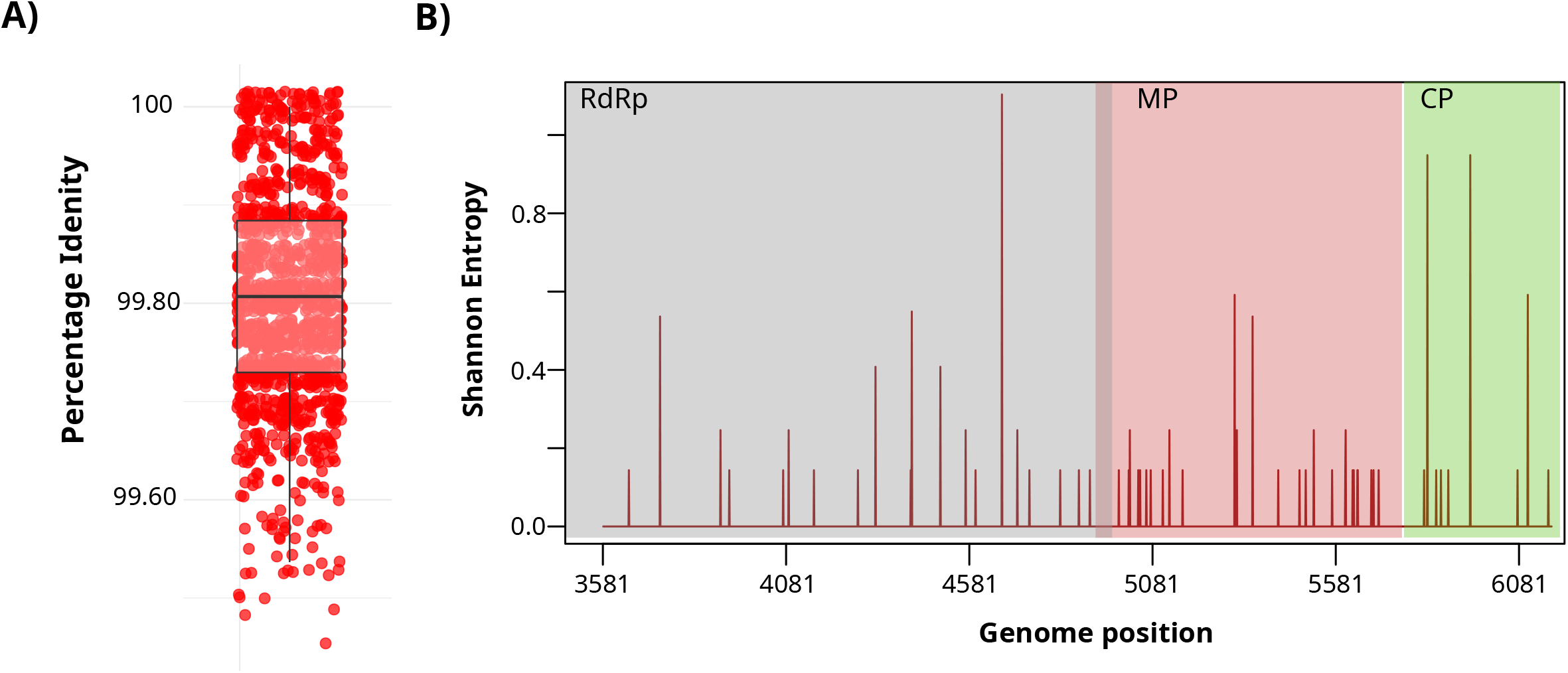
Genetic variability of tomato brown rugose fruit virus (ToBRFV) (Tobamovirus fructirugosum) sequences from Morocco. (A) Percentage of nucleotide identity among the different sequences from Morocco. (B) Genetic variability across the N terminus region of the RNA-dependent RNA polymerase (RdRp), the movement protein (MP), and the coat protein (CP) genes of ToBRFV using Shannon entropy.

We next investigated the selective pressures acting on each of these genes. No evidence of episodic diversifying selection was detected in entire genes, nor in single codons (Table S1, S2, and S3). However, we identified sites with nonsynonymous substitutions. A guanine was substituted by adenine in the site 5,304 of the genome within the MP gene with a variant frequency of 14.3% (p < 0.001), changing the codon from GAA to AAA; this substitution changed the amino acid from glutamic acid (negatively charged) to lysine (positively charged). Two non-synonymous substitutions were identified in the CP gene, one in position 5,947 substituting a thymine into adenine with a variant frequency of 36.7% resulting in the amino acid change from aspartic acid (codon GAU) to glutamic acid (codon GAA) (p < 0.001). Another mutation was detected in position 6,104 substituting the guanine into adenine with a variant frequency of 14.3% resulting in the amino acid change from valine (codon GUA) to isoleucine (codon AUA) (p < 0.001).

### Relationship among the ToBRFV sequences in Morocco and the rest of the world

We explored the relationship between the different ToBRFV sequences using haplotype network construction. Our results revealed 27 haplotypes clustered into four main groups (i to iv in Figure 3). Groups (i) and (ii) are central in the network, while groups (iii) and (iv) were located on the edges of the network. Groups (i) and (ii) exhibited predominant haplotypes connected to several variants by one to four mutations, while groups (iii) and (iv) are separated from the main clusters by intermediate haplotypes (Figure 3).

**Figure 3.**
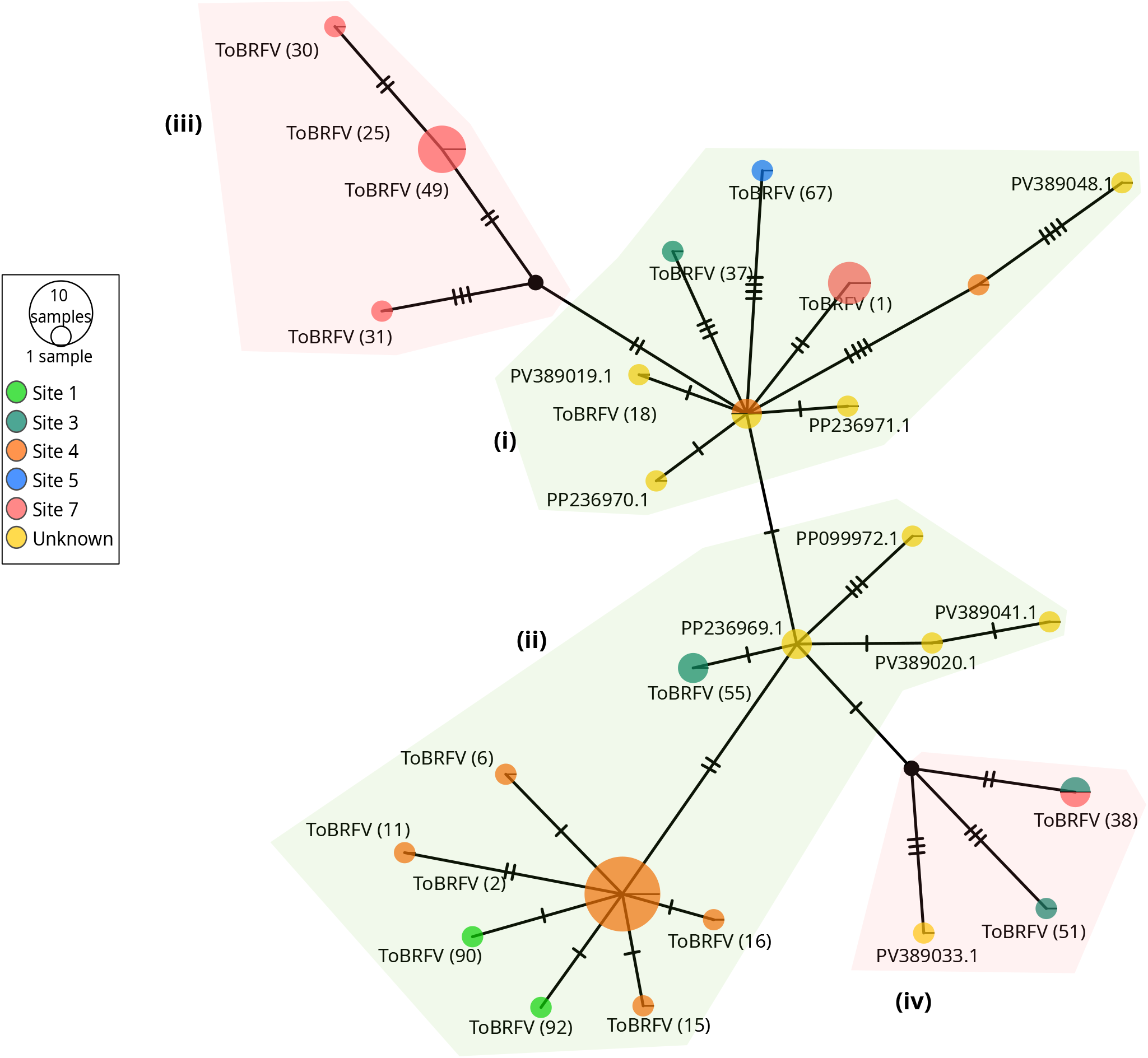
Relationships among the tomato brown rugose fruit virus (ToBRFV) (Tobamovirus fructirugosum) sequences from Morocco using “Median-joining” haplotype network. Different clusters or groups are highlighted in light green and light red.

The distribution of haplotypes across the different sites showed that several haplotypes were shared among the different locations, although no *sensu stricto* clustering was observed by the site. However, some subclusters were composed predominantly of samples from the same site, suggesting partial structuring within the population (Figure 3).

Next, we examined the relationships between ToBRFV sequences from Morocco and the rest of the world (Figure 4, Figure S1). We observed the clustering of the ToBRFV isolates into distinct clades. Sequences from Morocco positioned in different clusters. A major cluster comprised the majority of the sequences from Morocco. This cluster included sequences from Senegal, Netherland, and the UK. This cluster seems to have been emerged from sequences from Jordan and Israel. Two additional sequences were clustered in a group comprising different isolates which does not seem to be well differentiated. This group comprised sequences from all the continents which were separated by a small number of mutations. We also noticed that some clusters comprised sequences from China, others comprising sequences from The Netherlands. Sequences from The Netherlands appeared to be present in almost all the clusters.

**Figure 4.**
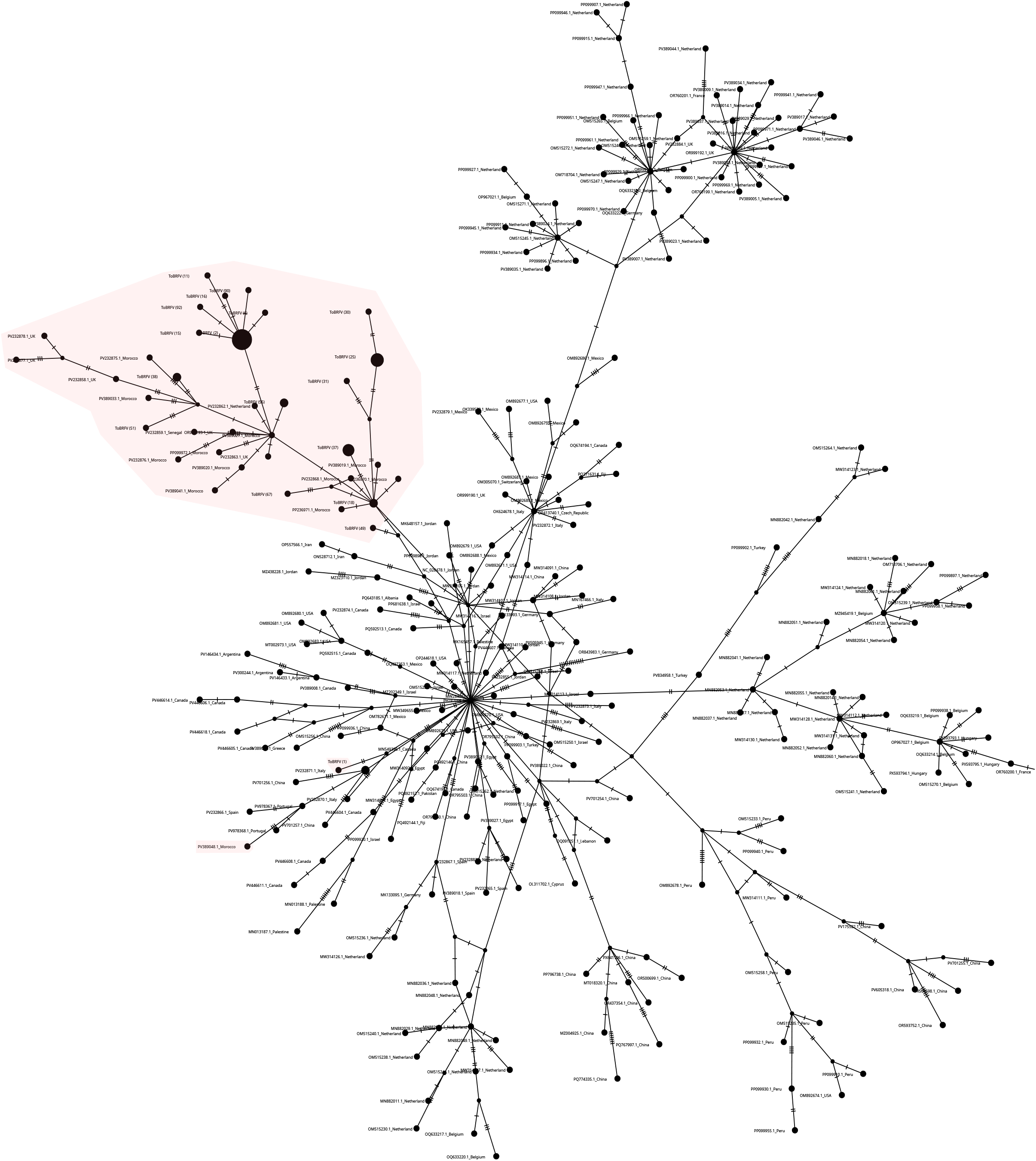
Relationships among the tomato brown rugose fruit virus (ToBRFV) (Tobamovirus fructirugosum) sequences from Morocco and the world using “Median-joining” haplotype network. ToBRFV sequences from Morocco are highlighted in light red. A network with higher resolution was provided in the supplementary material as Figure S1.

The overall network showed limited divergence among isolates, since haplotypes were connected by one to thirteen mutations, consistent with the low genetic variability of the virus. Clustering based on geographic origin was observed for the sequences from China, and for others from The Netherlands and Belgium, still the ones from different countries were intermixed throughout the network.

### Genetic diversity analyses of the movement protein of ToBRFV

Mutations in the MP help ToBRFV to evade *Tm-2*^*2*^ mediated resistance (Hak and Spegelman, 2021, Zisi et al., 2024, Ma et al., 2026). Therefore, we further studied the variability of the MP using globally available sequences (Figure 5). While this protein is well conserved, the variability observed was unevenly distributed across the protein (Figure 5A). The C-terminus, comprising the acidic and basic regions, exhibited higher variability than the N-terminus, which comprises the I and II domains. The E132K mutation observed in some ToBRFV sequences from Morocco was located in a conserved region within the domain II, closer to the N125, K129, and A134, which are critical residues for the function of the MP (Figure 5A and 5B).

**Figure 5.**
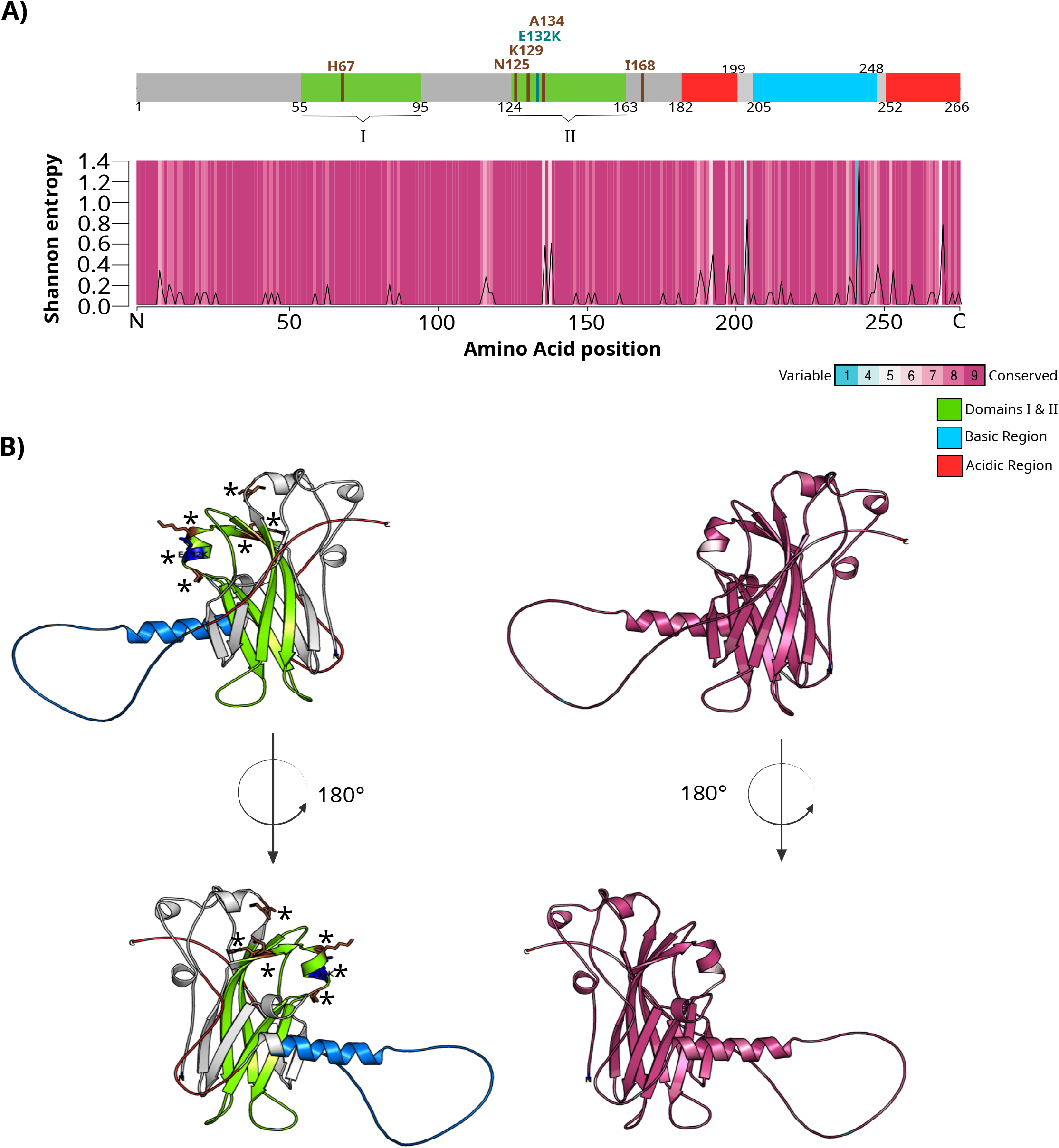
Genetic variability across the movement proteins (MP) of tomato brown rugose fruit virus (ToBRFV) (Tobamovirus fructirugosum) sequences using global sequences and ones generated from this study. Schematic representation of the different domains within the MP and the variability across them using the primary structure (A) and the quaternary structure (B). Important amino acid for the function of the MP are highlighted in brown in the primary and the quaternary structures (Ma et al., 2026). The MP substitution reported from sequences in this study, is highlighted in deep blue-green (Teal) in the primary structure (3A) and in blue in the quaternary structure (3B). Asterisks were added to the quaternary structure to facilitate the visuazilation of the important amino acids.

## Discussion

In this work, we used tomato fruits from local markets as sentinels to explore the virus genetic diversity in Morocco. The use of sentinels as a sampling strategy combined with molecular tools was shown to be efficient in documenting the virosphere and disease emergence in different ecosystems and species (Blondet et al., 2025).

The overall genetic diversity of the virus is low, which is consistent with the global situation and its recent introduction to the country since official reports date back to 2021. For this reason, we used haplotype network construction to depict their relationship. The network revealed a structured population composed of 27 haplotypes clustered around two major nodes. These central haplotypes probably represent ancestral variants from which the current ToBRFV population in Morocco has diversified. The limited number of mutational steps connecting the haplotypes (one to four mutations) suggests relatively recent diversification. This pattern supports a scenario of population expansion, which is common for emerging viruses (Holmes, 2009; Pybus and Rambaut, 2009). The presence of intermediate nodes may reflect unsampled haplotypes within the population or, alternatively, multiple independent introduction events of the virus to Morocco. Moreover, haplotype network analyses, using global sequences, showed a clustering of the Moroccan variants distinct from worldwide sequences, suggesting that geographical factors combined with agricultural practices may contribute to structuring the ToBRFV population from Morocco, playing a key role in shaping its evolutionary trajectory. The Moroccan cluster emerged from sequences from Jordan and Israel, suggesting a probable introduction from these countries. This cluster comprised sequences from the UK, Senegal, and The Netherland, suggesting probable dissemination of the Moroccan variants. The presence of two sequences, emerging from the ones from Italy, suggests a probable second route of introduction to the country. The low prevalence of these last variants may probably indicate their lack of competitiveness with the major ones observed in this study, or a lack of deep sampling to reveal their real prevalence across the country. Sequences from The Netherlands and Belgium were dispatched among the different clusters, reflecting higher variability and probably suggesting multiple introductions of the virus to these countries and/or their role in its spread to other countries through seed industry.

While previous studies have shown that the MP gene of ToBRFV exhibits the highest genetic variability at the global scale, suggesting its implication in virus evolutionary adaptation (Ghorbani et al., 2024), our results show that the CP exhibited almost 1.7 times more diversity than the MP and the partial RdRp sequences. Despite CP variability, no episodic diversifying selection was identified, and the detected amino acid substitutions were conservative (D78E and V131I), suggesting that these variations are tolerated without strongly affecting function, rather than being driven by adaptive diversification.

On the other hand, the MP substitution (E132K) involved a polarity change and may probably have greater functional relevance despite its lower frequency. The MP of tobamoviruses contains two highly hydrophobic regions that are predicted to span membranes (Brill et al., 2000). The MP causes transient aggregation of host endoplasmic reticulum, which apparently plays a role in formation of cytoplasmic bodies where tobamoviruses replicate (Heinlein et al., 1998; Más and Beachy 1999). The MP is the primary determinant of resistance to tobamoviruses in hosts carrying the *Tm*-2^2^ resistance gene. ToBRFV was able to overcome that resistance present in the majority of commercial tomato cultivars. Mutations in the MP help ToBRFV to evade *Tm*-2^2^ mediated resistance by attenuating MP–*Tm*-2^2^ interaction strength, thereby avoiding *Tm*-2^2^ self-association and downstream immune activation (Ma et al., 2026). While mutations in the MP can abolish the cell-to-cell and systemic movement (Ma et al., 2026; Figure 3), a recent study showed that a single mutation in ToBRFV breaks virus-specific resistance in new resistant tomato cultivars (Zisi et al., 2024). The mutation was located within the MP substituting a (T) to (G), leading to amino acid change N82K in the MP of the resistance breaking isolate (Zisi et al., 2024). We found that the MP of ToBRFV is highly conserved across the global sequences, still the C-terminal comprising the acidic and basic regions are more variable. The mutation found in some of the Moroccan sequences fall in a conserved residue within the domain II. This residue does not fall within the ones reported to inhibit the MP function, leading probably to adaptive diversification. Further biological assays should be carried out to assess whether the current mutation in these variants provide functional adaptation.

In conclusion, our study establishes the basis for tracking the evolutionary dynamics of ToBRFV in Morocco. Future work is needed to characterize the biological relevance of the observed pattern among the Moroccan isolates with respect to host responses.

## Acknowledgments

We thank Fatima Valle (CEBAS-CSIC) for technical assistance.

## Data availability

Virus sequences were deposited in the GenBank database under the following accession numbers: ToBRFV Tom1 (PZ370457), ToBRFV Tom2 (PZ370458),ToBRFV Tom3 (PZ370459),ToBRFV Tom5 (PZ370460),ToBRFV Tom6 (PZ370461),ToBRFV Tom7 (PZ370462),ToBRFV Tom9 (PZ370463),ToBRFV Tom10 (PZ370464),ToBRFV Tom11 (PZ370465),ToBRFV Tom12 (PZ370466),ToBRFV Tom13 (PZ370467),ToBRFV Tom15 (PZ370468),ToBRFV Tom16 (PZ370469),ToBRFV Tom18 (PZ370470),ToBRFV Tom19 (PZ370471),ToBRFV Tom20 (PZ370472),ToBRFV Tom21 (PZ370473),ToBRFV Tom23 (PZ370474),ToBRFV Tom25 (PZ370475),ToBRFV Tom27 (PZ370476),ToBRFV Tom28 (PZ370477),ToBRFV Tom30 (PZ370478),ToBRFV Tom31 (PZ370479),ToBRFV Tom34 (PZ370480),ToBRFV Tom36 (PZ370481),ToBRFV Tom37 (PZ370482),ToBRFV Tom38 (PZ370483),ToBRFV Tom39 (PZ370484),ToBRFV Tom40 (PZ370485),ToBRFV Tom42 (PZ370486),ToBRFV Tom49 (PZ370487),ToBRFV Tom50 (PZ370488),ToBRFV Tom51 (PZ370489),ToBRFV Tom55 (PZ370490),ToBRFV Tom56 (PZ370491),ToBRFV Tom67 (PZ370492), ToBRFV Tom90 (PZ370493), ToBRFV Tom92 (PZ370494).

## Supplementary data

**Table S1**. Selective pressures acting on site-by-site codon on the partial RNA-dependent RNA polymerase gene.

**Table S2**. Selective pressures acting on site-by-site codon on the movement of protein gene.

**Table S3**. Selective pressures acting on site-by-site codon on the capsid protein gene.

**Figure S1**. Relationships among the tomato brown rugose fruit virus (ToBRFV) (*Tobamovirus fructirugosum*) sequences from Morocco and the world using “Median-joining” haplotype network. ToBRFV sequences from Morocco are highlighted in light red.

