## Supplementary material for "Genetic diversity of tomato brown rugose fruit virus in Morocco": Figure S1

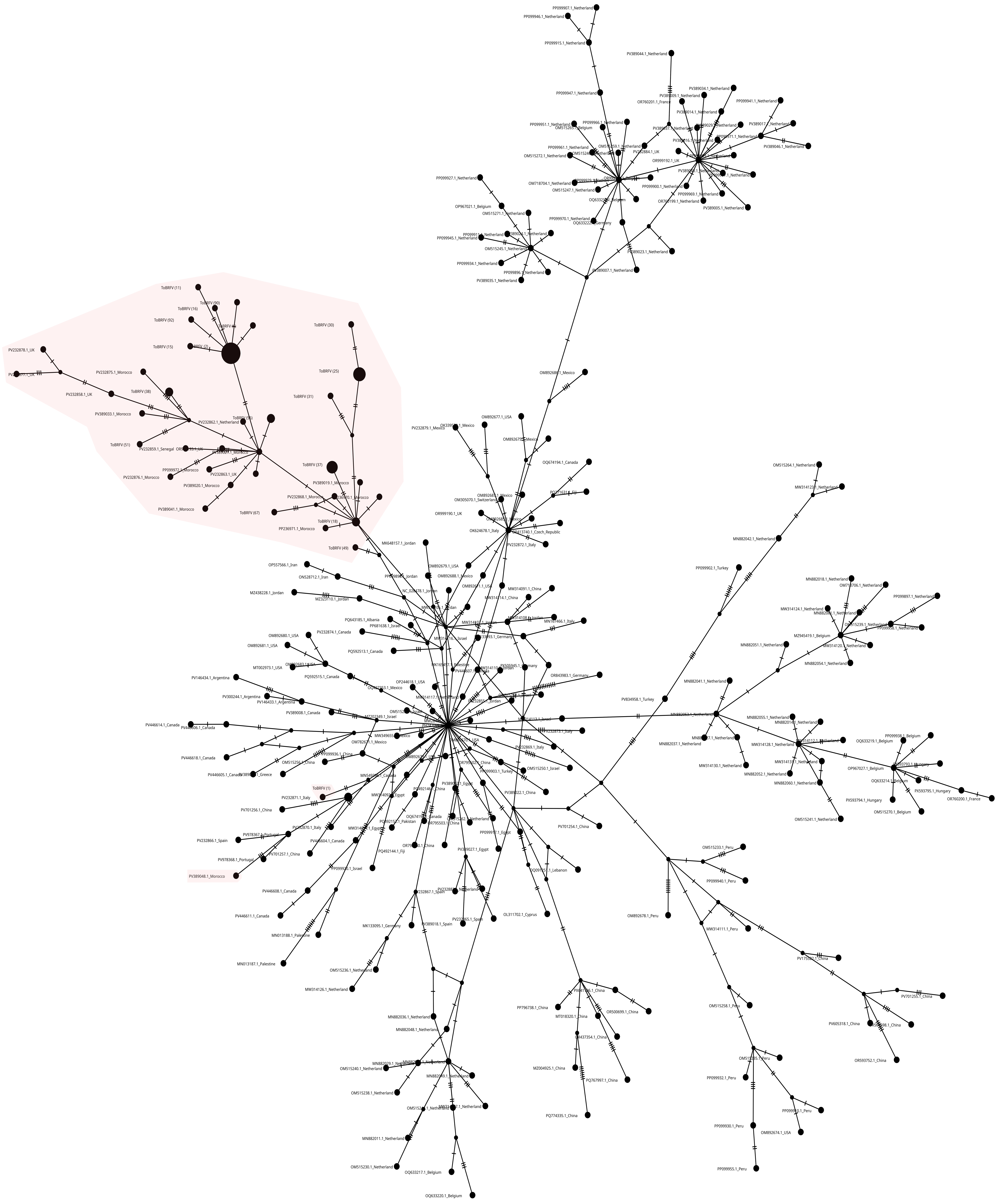

**Figure S1.** Relationships among the tomato brown rugose fruit virus (ToBRFV) (Tobamovirus fructirugosum) sequences from Morocco and the world using “Median-joining” haplotype network. ToBRFV sequences from Morocco  are highlighted in light red.
